# ReDis: efficient metagenomic profiling via assigning ambiguous reads

**DOI:** 10.1101/2023.08.29.555244

**Authors:** Chun Song, Zule Guo, Ju Gu, Yong Ren, Hao Guo, Junfeng Liu

**Affiliations:** State Key Laboratory of Translational Medicine and Innovative Drug Development, Jiangsu Simcere Diagnostics Co., Ltd., Nanjing, China; Nanjing Simcere Medical Laboratory Science Co., Ltd., Nanjing, China

## Abstract

**Summary:** Metagenomic profiling is one of the primary means of microbiome analysis, which includes classification of sequencing reads and quantification of their relative abundances. Although mang methods have been developed for metagenomic profiling, metagenomic profiling remains challenges on striking a delicate balance between accuracy and runtime as well as assigning the ambiguous reads. Here, we present a novel method, named ReDis, to overcome the above issues. ReDis combines Kraken2 with Minimap2 for aligning sequencing reads against a reference database with hundreds of gigabytes (GB) in size accurately within feasible time, and then uses a novel statistical model to assign the ambiguous reads for producing accurate abundance estimates. In contrast to the popular Kraken2+Bracken, ReDis improved the accuracy of abundance estimation on simulated reads from two highly similar genomes: *Escherichia coli* and *Shigella flexneri*.

## 1 Introduction

Metagenomic profiling aims to identify species contained in a metagenomic sample by classification of sequencing reads and quantification of their relative abundances and is one of the primary means of microbiome analysis. Currently, various metagenomic profilers that rely on reference database have been developed to classify metagenomic data and estimate taxon abundance profiles (Simon *et al*., 2019; Sun *et al*., 2021). There are two challenges for profiling metagenomic data. The first challenge is how to strike a delicate balance between accuracy and runtime for aligning sequencing reads against a reference database with hundreds of gigabytes (GB) in size. Alignment-based methods are highly accurate, yet computationally infeasible when the reference database reaches to hundreds of gigabytes (GB) in size (LaPierre *et al*., 2020). Although the k-mer-based approaches can provide a fast metagenomic sequence classification (Simon *et al*., 2019; Wood *et al*., 2019), they are typically not as accuracy as the alignment-based methods, especially in differentiating highly similar genomes (Sheka *et al*., 2021; Liao *et al*., 2022). The second challenge is how to count and assign the ambiguous reads that align equally well to more than one genome. There are two typical methods for counting the ambiguous reads, which respectively are Centrifuge (Kim *et al*., 2016) and Bracken (Lu *et al*., 2017). However, Centrifuge neglects the effect of the similarities of genomes in a reference database on counting the ambiguous reads. Although the similarities of genomes in a reference database have been considered by Bracken, the statistical model of Bracken can be only used to count the ambiguous reads in conjunction with Kraken (Wood and Salzberg, 2014) or Kraken2 (Wood *et al*., 2019). In addition, to the best of our knowledge, the tool that is able to assign the ambiguous reads has yet been found in literature.

In this study, we propose a novel method, named ReDis, to address common obstacles of metagenomic analyses. We first combine Kraken2 with Minimap2 (Li, 2018) for aligning metagenomic sequences in order to achieve a strong balance of accuracy and runtime. Then, we apply Bayes’ theorem to establish a statistical model for computing the probability that an arbitrary read belongs to one given genome. In the established statistical model, the influence of similarities of genomes in a reference database is described by unique mapping rate. And finally, based on the probability that an ambiguous read belongs to one given genome, we assign the ambiguous read to a specified genome through pseudo-random number generator. We demonstrate that ReDis can be effective to tackle challenges of metagenomic profiling by simulations.

## 2 Implementation

The input to ReDis is sequencing reads and a reference database. We adopt three strategies in ReDis for the balance between highly accurate alignment and runtime as well as performing efficient metagenomic profiling. The first strategy is the reference database pre-filtering by running alignment with Kraken2 (Fig. 1a). We use Kraken2, which is the fastest metagenomic classifier (Meyer *et al*., 2022), to align metagenomic sequences to the full reference database and select the reference genomes to which reads have been classified to build a sub-database (see Methods section “Sub-database construction” in Supplementary Materials). Then, we perform accurate alignment of reads to the sub-database in much less time using Minimap2 and construct the independent subsets (see Methods section “Independent subsets” in Supplementary Materials) (Fig. 1b). The second strategy is introducing unique mapping rate to establish the statistical model for computing the probability that the arbitrary read in each independent subset belongs to one given genome. The established statistical model is based on Bayes ’ theorem and the independent subsets are divided from alignment results from Minimap2 (see Methods section “Statistical model” in Supplementary Materials). The third strategy is using pseudo-random number generator to assign the arbitrary ambiguous read to a specified genome. We compute the probability that an ambiguous read belongs to one given genome and decide whether the ambiguous read is assigned to the given genome according to the number produced by pseudo-random number generator (see Methods section “Assigning ambiguous reads” in Supplementary Materials). Finally, ReDis outputs a profile that reports relative sequence abundance by combining alignment results of reads and assignment results of ambiguous reads (see Methods section “Abundance analysis” in Supplementary Materials) (Fig. 1c).

**Fig. 1.**
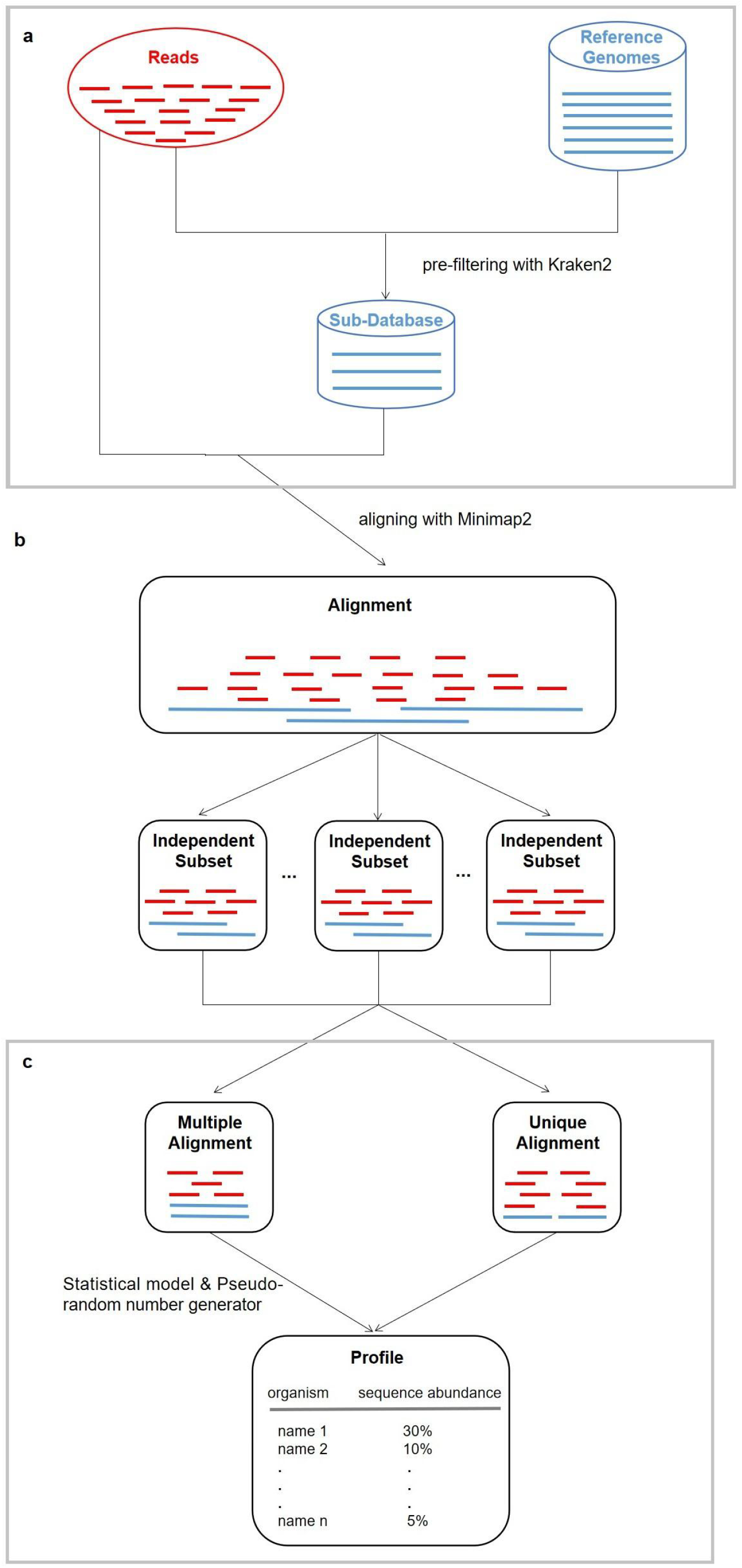
ReDis overview. (A) Building a sub-database. The input to ReDis is sequencing reads and a reference database. ReDis then filters the reference database by running alignment with Kraken2. (B) Constructing independent subsets. ReDis performs accurate alignment of reads to the sub-database using Minimap2 and constructs the independent subsets. (C) Assigning ambiguous reads with a statistical model and pseudo-random number generator. ReDis assigns the arbitrary ambiguous read to a specified genome and outputs the relative sequence abundance.

## 3 Results and discussion

We applied ReDis to differentiate highly similar species on genome, such as *Escherichia coli* and *Shigella flexneri*, which are well over 99% identical (Jin *et al*., 2002). For the five simulated microbial communities with only two species: *Escherichia coli* and *Shigella flexneri* (see Methods section “Simulation of sequencing reads” in Supplementary Materials), the ratio of *Escherichia coli* to *Shigella flexneri* is 1:1 and the numbers of reads of each species are 20, 100, 500, 2,500, and 12,500 respectively. Compared with Kraken2+Bracken, which has been applied in thousands of published microbiome studies (Sun *et al*., 2021), the simulation results showed that ReDis performed better in the read number estimation except for the estimation of *Escherichia coli* on the simulated dataset with 200 reads (Supplementary Table S1).

The outperformance of ReDis is because ReDis can assign the ambiguous reads to specific genomes by computing the probability that each ambiguous read derives from each aligned genome. On the five simulated datasets, the average proportions of correct reads assigned to *Escherichia coli* and *Shigella flexneri* by ReDis are 0.75 and 0.72, respectively (Supplementary Table S2). When only considering the wrong reads assigned to other organisms (not including *Escherichia coli* and *Shigella flexneri*), the average proportions of wrong reads are 0.04 and 0.02 respectively (Supplementary Table S2). In addtion, for the ambiguous reads identified by Kraken2 on the five simulated datasets, the re-distribution of the ambiguous reads by ReDis is more closer to the true distribution than the re-distribution of the ambiguous reads by Bracken (Supplementary Table S3). All of this show that ReDis can assign the ambiguous reads effectively.

We run Minimap2 on the five simulated datasets and the average runtime was more than 34 minutes (Supplementary Table S4). Due to constructing sub-database, ReDis took less than 6 minutes in all cases (Supplementary Table S4). Although ReDis was slightly slower than Kraken2+Bracken that took about 4 minutes (Supplementary Table S4), ReDis is able to assign the ambiguous reads to specific genomes and produce good abundance estimates. Therefore, ReDis achieves a delicate balance between accurate metagenomic profiling and runtime.

## 4 Conclusion

In this study, we developed ReDis that is a novel approach for classification of metagenomic data and estimation of taxonomic profiles. ReDis improves the performance of abundance estimation, which is critical for precision-recall analysis. Two factors drive ReDis’s high performance. One factor is that ReDis’s sub-database construction step substantially quickens downstream alignment and increases the precision of alignment due to the removal of reference genomes of organisms that cannot reasonably be contained in the sample by Kraken2. The other factor is that ReDis ’s novel assigning ambiguous reads step significantly raises the accuracy of abundance estimation of the organism with many multi-mapped reads by establishing the statistical model including the unique mapping rate. In addition, ReDis is able to estimate the sequence abundance at any taxonomic rank. For these reasons, we anticipate that ReDis will be widely applied for microbiome analysis.

