## Supplemental Table 1-4 for "ReDis: efficient metagenomic profiling via assigning ambiguous reads"

### 1. Methods

#### Reference Database construction

ReDis supports custom reference database, which is created by the end user. In this study, we use the most popular reference database, RefSeq complete genomes (RefSeq CG), from bacterial, fungi, and viral kingdoms. The final database consists of 681 families totaling 90 GB in size. We use ascp to download all RefSeq CG and then filter out unwanted taxa such as animals and plants.

#### Sub-database construction

We use Kraken2 (2.1.2) to align reads of the input sample to the full reference database via the default parameters. Based on the taxonomic tree outputted by Kraken2, we select the genomes from the full reference database at each taxonomic rank from strain to family to construct sub-database. For the strain-level taxa, we select the genomes from the full reference database, which are on the strain leaves in the taxonomic tree. For the species-level taxa, we select the reference genomes that are on the species nodes in the taxonomic tree or their strains. At other two taxonomic levels (genus or family), the number of reads assigned to nodes in the taxonomic tree should cross the given threshold. We specify that the thresholds of genus and family are 100 and 1,000 respectively. For one genus or family node in the taxonomic tree, to which the number of reads assigned is over the given threshold, we select the reference genomes that are on its child species or strains. Based on the much smaller sub-database, we can perform accurate alignment of reads of the input sample in a reasonable amount of time using Minimap2 (2.22).

#### Independent subsets

We divide the reads of input sample into several independent subsets according to the alignment results outputted by Minimap2. The independent subset (denoted as $R_i$) is defined as follows: (1)

$R_1 \cup R_2 \cup \cdots \cup R_m = R$ ; (2) $R_i \cap R_j = \Phi \quad i \neq j \quad i, j = 1,2,\cdots,m$ . $R$ represents all reads in the input sample and $m$ is the number of independent subsets. If the intersection of $R_i$ and $R_j$ is null, they do not share any read as well as any reference genome to which the reads in $R_i$ or $R_j$ can be aligned. Furthermore, the independent subset $R_i$ cannot be subdivided following the above definition. On each independent subset, we can compute the probability that the arbitrary read belongs to one given genome by constructing statistical model.

### Statistical model

We define the following statistical model with Bayes' theorem for computing the probability that a read in the independent subset $R_j$ emanates from genome $S_i$: $P(S_i|R_j) = \frac{P(R_j|S_i)P(S_i)}{P(R_j)}$, where $P(R_j|S_i)$ is the probability that a read from genome $S_i$ is classified to $R_j$, $P(S_i)$ is the probability that a read in the input sample belongs to genome $S_i$, and $P(R_j)$ is the probability that a read in the input sample is classified to $R_j$. According to the definition of independent subset, $P(R_j|S_i)$ is 1 if there exist reads in $R_j$ that can be aligned to genome $S_i$ and 0 otherwise. Let $P(S_i)$ be expressed as follows: $P(S_i) = \frac{\frac{N_i}{K_i}}{\sum_{l=1}^{n} \frac{N_l}{K_l}}$, where $N_i$ is the number of the reads in the input sample that are uniquely aligned to genome $S_i$, $K_i$ is the unique mapping rate of genome $S_i$, and $n$ is the number of genomes to which all reads in the input sample can be aligned. Based on the law of total probability, $P(R_j)$ can be expressed as follows: $P(R_j) = \sum_{i=1}^{n} P(R_j|S_i)P(S_i)$.

The unique mapping rate of genome $S_i$ is the proportion that nucleotide bases on the reads uniquely aligned to genome $S_i$ is to the total nucleotide bases of genome $S_i$. In order to count the nucleotide bases on the unique reads aligned to genome $S_i$ of length $L_i$, we generate ($L_i$-$r$+1) reads across the entire genome by a sliding window of length $r$ and align the generated reads to sub-database using Minimap2. The length $r$ is the average length of reads in the input sample. For example, if $L_i$=1 Mbp, $r$=100 bp, and only 3,000 reads that cover 250,000 nucleotide bases of genome $S_i$ are unique, then $K_i$=0.25.

### Assigning ambiguous reads

For a given ambiguous read in the independent subset $R_j$, if it was aligned equally well to genome $S_1$, genome $S_2$, and genome $S_3$, then we can compute the probability that it derives from genome $S_i$ ($i$=1,2,3) as follows: $q_i = \frac{P(S_i|R_j)(1-K_i)}{\sum_{l=1}^{3} P(S_l|R_j)(1-K_l)}$ $i=1,2,3$. According to the computed probabilities, we break the interval (0,1) into three segments of lengths $q_1$, $q_2$, and $q_3$. We then draw a random number $\mu$ from the uniform distribution $U(0,1)$ by pseudo-random number generator which is a mathematical algorithm that produces a sequence of numbers between 0 and 1 that appear random. At last, we assign the given ambiguous read depending on which segment $\mu$ falls into. For example, if $q_1$=0.2, $q_2$=0.3, and $q_3$=0.5, then the interval (0,1) will be divided into three segments: (0,0.2], (0.2,0.5], and (0.5,1). When the generated random number $\mu$ falls into the interval (0.5,1), such as $\mu$=0.7, the given ambiguous read should be assigned to genome $S_3$.

### Abundance analysis

ReDis can perform abundance analysis at any taxonomic rank, e.g., strain, species, genus, or higher taxonomic level. For a given taxon, ReDis re-distributes the reads assigned to each genome included in the given taxon upward to it and counts the number of the re-distributed reads. The abundance of the given taxon is calculated as the proportion of reads re-distributed to it out of the total number of reads in the input sample. For example, if the given taxon $k$ includes three genomes and the numbers of the reads assigned to each genome are $n_1$, $n_2$, and $n_3$, respectively, then the abundance of $k$ is $(n_1+n_2+n_3)/n$ where $n$ is the number of total reads in the input sample. ReDis outputs sequence abundance, but not taxonomic abundance which is denoted as the number of genomes of a given taxon relative to the total number of genomes detected. The taxonomic abundance of a given taxon $k$ can be expressed as follows: $\frac{N_k/L_k}{\sum_{i=1}^{m} N_i/L_i}$, where $N_k$ is the number of reads re-distributed to taxon $k$, $L_k$ is the average length of genomes included in taxon $k$, and $m$ is the number of identified taxa in the input sample. It is notable, however, that sequence abundance and taxonomic abundance are not related by universal algebraic relation (Sun *et al.,* 2021).

### Simulation of sequencing reads

We downloaded the reference genomes of *Escherichia coli* and *Shigella flexneri* from NCBI: GCF_000005845.2_ASM584v2_genomic.fna, GCF_000006925.2_ASM692v2_genomic.fna. Then, we used ART (Huang *et al.,* 2012) with default parameters to simulate single-end sequencing reads based on HS2500 platform. We generated five simulation datasets: S-1, S-2, S-3, S-4, and S-5, in which the length of each read is 75bp and the ratio of *Escherichia coli* to *Shigella flexneri* is 1:1. In five simulation datasets, the numbers of reads are 40, 200, 1,000, 5,000, and 25,000.

### 2. Running ReDis and Kraken2+Bracken

#### ReDis

```
./script/redis_v1 \
        -i redis_input_list.txt \
        -o ./res_redis \
        -d ./db_redis \
        -m /apps/bin/minimap2 \
        -l 75 \
        -t 50 \
        -r 1
```

#### Kraken2+Bracken

Kraken2:

```
/kraken2_braken/kraken2/kraken2 \
        --thread 8 \
        --db /krakendb \
        --report /sim_40.kraken.report \
        --output /sim_40.kraken.output \
        /sim_40.fa
```

Bracken:

```
/kraken2_braken/Bracken/bracken \
        -d /krakendb \
        -i /sim_40.kraken.report \
        -r 75 \
        -l S \
        -t 1 \
        -o
```

### 3. Simulation Results

**Table S1.** The sequence abundance estimations on the simulation datasets.

| Simulation Dataset | Kraken2+Bracken | | ReDis | |
|---|---|---|---|---|
| | *Escherichia coli* | *Shigella flexneri* | *Escherichia coli* | *Shigella flexneri* |
| S-1 | 0.25 | 0.725 | 0.475 | 0.45 |
| S-2 | 0.51 | 0.48 | 0.455 | 0.49 |
| S-3 | 0.618 | 0.378 | 0.52 | 0.432 |
| S-4 | 0.578 | 0.418 | 0.496 | 0.43 |
| S-5 | 0.58 | 0.415 | 0.472 | 0.425 |

Note. Each simulation dataset only includes two species: *Escherichia coli* and *Shigella flexneri,* with the ratio of 1:1. The numbers of reads of five simulation datasets are 40, 200, 1,000, 5,000, and 25,000 respectively.

**Table S2.** The details of assigning the simulated reads by ReDis.

| Simulation Dataset | *Escherichia coli* | | | *Shigella flexneri* | | |
|---|---|---|---|---|---|---|
| | Correct assigned reads | Wrong assigned reads (to *Shigella flexneri*) | Wrong assigned reads (to other organisms) | Correct assigned reads | Wrong assigned reads (to *Escherichia coli*) | Wrong assigned reads (to other organisms) |
| S-1 | 14 | 4 | 0 | 14 | 5 | 0 |
| S-2 | 69 | 25 | 3 | 73 | 22 | 1 |
| S-3 | 377 | 93 | 10 | 339 | 143 | 2 |
| S-4 | 1798 | 476 | 134 | 1674 | 684 | 35 |
| S-5 | 8592 | 2337 | 1150 | 8296 | 3212 | 656 |

Note. Each simulation dataset only includes two species: *Escherichia coli* and *Shigella flexneri*, with the ratio of 1:1. The numbers of reads of five simulation datasets are 40, 200, 1,000, 5,000, and 25,000 respectively.

**Table S3.** The re-distribution of ambiguous reads identified by Kraken2 on the simulation datasets.

| Simulation Dataset | The true distribution of ambiguous reads identified by Kraken2 | | The re-distribution of ambiguous reads by Bracken | | The re-distribution of ambiguous reads by ReDis | |
|---|---|---|---|---|---|---|
| | *Escherichia coli* | *Shigella flexneri* | *Escherichia coli* | *Shigella flexneri* | *Escherichia coli* | *Shigella flexneri* |
| S-1 | 18 | 14 | 8 | 23 | 17 | 13 |

| | | | | | | |
|---|---|---|---|---|---|---|
| S-2 | 83 | 63 | 86 | 59 | 77 | 60 |
| S-3 | 370 | 379 | 489 | 258 | 417 | 300 |
| S-4 | 1977 | 1801 | 2373 | 1396 | 2061 | 1444 |
| S-5 | 9906 | 9176 | 11945 | 7086 | 9724 | 7317 |

Note. Each simulation dataset only includes two species: *Escherichia coli* and *Shigella flexneri*, with the ratio of 1:1. The numbers of reads of five simulation datasets are 40, 200, 1,000, 5,000, and 25,000 respectively.

**Table S4.** The wall clock running time on simulation datasets.

| Simulation Dataset | Wall clock running time (minutes) | | |
|---|---|---|---|
| | Kraken2+Bracken | ReDis | Minimap2 |
| S-1 | 3:52 | 5:01 | 34:54 |
| S-2 | 4:14 | 5:19 | 34:42 |
| S-3 | 4:01 | 5:07 | 34:40 |
| S-4 | 3:56 | 5:02 | 34:52 |
| S-5 | 3:41 | 5:48 | 35:18 |

Note. Each simulation dataset only includes two species: *Escherichia coli* and *Shigella flexneri*, with the ratio of 1:1. The numbers of reads of four simulation datasets are 40, 200, 1,000, 5,000, and 25,000 respectively. The reference database consists of 681 families totaling 90 GB in size. Kraken was run with 8 threads. Bracken was run with 1 thread. Redis and Minimap2 were run with 20 threads.
